# The thalamus regulates retinoic acid signaling and development of parvalbumin interneurons in postnatal mouse prefrontal cortex

**DOI:** 10.1101/272427

**Authors:** Rachel Larsen, Alatheia Proue, Earl Parker Scott, Matthew Christiansen, Yasushi Nakagawa

## Abstract

Abnormal development of GABAergic interneurons in the prefrontal cortex (PFC) is implicated in a number of psychiatric disorders. Yet, developmental mechanisms for these neurons are poorly understood. Here we show that the retinoic acid-degrading enzyme CYP26B1 is temporally expressed specifically in postnatal frontal cortex in mice, and its genetic deletion results in an increased density of parvalbumin (PV)-expressing interneurons in medial PFC during postnatal development. Furthermore, initiation of *Cyp26b1* expression in neonatal PFC depends on the connections between the thalamus and the neocortex. Thus, the thalamus has a postnatal role in regulating PV neuron development in PFC by inducing *Cyp26b1* and thereby restricting retinoic acid signaling. Prenatally, the lack of thalamic input causes an aberrant radial distribution of medial ganglionic eminence-derived interneurons throughout the cortex. Therefore, the thalamus controls PV neuron development in PFC both by region-specific and cortex-wide mechanisms.

## Introduction

The prefrontal cortex (PFC) has major connections with several brain regions including the thalamus, amygdala, hippocampus and striatum, and integrates many modalities of information to execute higher functions such as goal-oriented behaviors, social interactions and emotion. Aberrant development of the PFC has been linked to schizophrenia, autism spectrum disorders, attention deficit hyperactivity disorders (ADHD), depression and bipolar disorders^1^. More specifically, developmental trajectories of GABAergic interneurons in the PFC, particularly those expressing the calcium-binding protein parvalbumin (PV), are impaired in both human patients and animal models of these disorders^2–7^. Therefore, determining the developmental mechanisms of PFC interneurons is important for understanding the disease pathophysiology.

Development of cortical interneurons is regulated by both intrinsic and extrinsic mechanisms^8–11^. One key extrinsic cue is the input from the thalamus, which regulates migration and maturation of cortical interneurons both prenatally and postnatally in mice^12–15^. However, most studies addressing the role of the thalamic input in interneuron development have been performed on primary visual or somatosensory cortex, leaving the mechanisms in PFC understudied. The delayed maturation of PFC interneurons compared with other cortical areas^16–18^ and the distinct set of thalamic nuclei connected with the PFC^19,20^ suggest the presence of unique extrinsic regulatory mechanisms for interneuron development in the PFC.

One candidate molecule that may play a role in postnatal development of the PFC is retinoic acid (RA), a small molecule derived from vitamin A. RA is critical for many important aspects of brain development, ranging from rostrocaudal patterning of the hindbrain and spinal cord to synaptic plasticity^21–25^. The RA-degrading enzyme, CYP26B1, is crucial in embryonic development in vertebrates^26–28^. In the mouse neocortex, *Cyp26b1* is expressed in the deep layer of the frontal cortex (Allen Brain Atlas). In contrast, *Aldh1a3*, which encodes an RA-synthesizing enzyme, is expressed in the superficial layer of the medial PFC during postnatal development^29^. The already established role of RA in embryonic brain development and the unique opposing locations of *Cyp26b1* and *Aldh1a3* expression imply that the balance between degradation and production of RA might play an unexplored role in postnatal development of medial PFC.

In this study, we first demonstrated that in medial PFC, *Cyp26b1* is transiently expressed in layer 6 cells from postnatal day 2 (P2) to P21, a period that overlaps with the early maturation of interneurons in the PFC, and that a significant subpopulation of PV interneurons are the main cell type that responds to RA. These results led us to hypothesize that RA signaling regulated by *Cyp26b1* plays a role in controlling the development of PV interneurons in the PFC. To test this, we generated layer 6- and frontal cortex-specific *Cyp26b1* mutant mice and found that they had an increased density of PV-expressing neurons in deep layers of medial PFC at P14 and P21. We further demonstrated that the postnatal expression of *Cyp26b1* in medial PFC is dependent on the connections between the thalamus and PFC. Thus, the thalamus has a postnatal role in regulating the development of PV neurons via inducing the expression of the RA-degrading enzyme CYP26B1 in frontal cortex. Additionally, we found that the thalamus is also required for the radial allocation of PFC interneurons during embryogenesis. These results demonstrate that the thalamus plays multiple crucial roles in interneuron development in the PFC first by controlling their laminar positioning during embryonic stages and then by restricting the maturation of PV neurons by inducing a retinoic acid-degrading enzyme.

## Results

### *Cyp26b1,*a gene encoding a retinoic acid degrading enzyme, is expressed in a spatially and temporally dynamic pattern in postnatal mouse PFC

In order to identify genes that are enriched in developing PFC, we screened the Anatomic Gene Expression Atlas (AGEA)^30^. The database showed that *Cyp26b1*, a gene that encodes a retinoic acid (RA)-degrading enzyme that belonged to the cytochrome P450 family 26 (CYP26) proteins, is strongly expressed in the frontal cortex at postnatal day 14 (P14), including the deep layer of medial and ventral PFC as well as the middle layer of lateral cortex extending into the agranular insula. We therefore examined the developmental expression patterns of *Cyp26b1* in more detail by in situ hybridization. Prenatal cortex did not show detectable *Cyp26b1* expression (Fig.1A). At P0, lateral frontal cortex, but not medial PFC, started to show a robust signal (Fig.1B). At P2, strong expression of *Cyp26b1* was detected in medial PFC, as well as in the dorsolateral frontal cortex including the motor area (Fig.1C). Comparison with the established marker of layer 6 neurons, *Synaptotagmin 6 (Syt6)*, showed that *Cyp26b1* is expressed in layer 6 of medial and ventral PFC (compare Fig.1D and M). Not only was the expression of *Cyp26b1* spatially restricted, it was also temporally dynamic; in medial PFC, *Cyp26b1* was strong at P8 (Fig.1D) and P14 (Fig.5C), but by P21, it was much weaker compared with ventral and lateral regions (Fig.1E). By P35, there was no detectable expression of *Cyp26b1* in medial PFC (Fig.1F).

**Fig.1.**
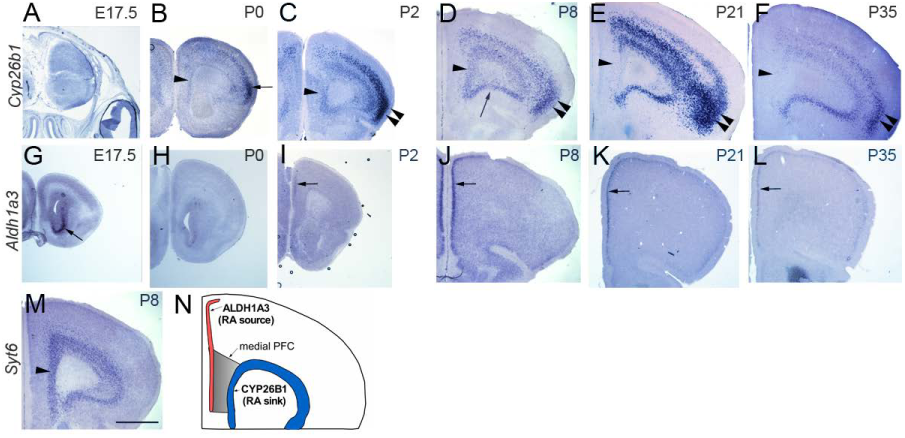
**Spatiotemporally regulated expression of *Cyp26b1* and *Aldh1a3* in the PFC** In situ hybridization of frontal sections through PFC at various stages is shown. **A-F.** *Cyp26b1* expression in frontal cortex. At E17.5, *Cyp26b1* is not detected (A). At P0, only a weak expression is seen in medial PFC (B, arrowhead). At P2, *Cyp26b1* starts to be detected clearly in medial PFC (C, arrowhead). Expression in lateral cortex, especially agranular insula in more superficial layer (B, arrow) is strong, which continues into later stages (C-F, double arrowheads). At P8, expression in medial (arrowhead) and ventral (arrow) PFC is strong (D). At P21, expression of *Cyp26b1* is reduced in medial PFC (E, arrowhead) and is almost undetectable by P35 (F, arrowhead). **G-L.** *Aldh1a3* expression in frontal cortex. At E17.5, expression is detected in LGE (G, arrow) but not in PFC. At P2, clear expression is detected in the superficial layer of medial PFC (I, arrow). The expression continues into P8, P21 and P35 (J-L, arrow). **M.** *Synaptotagmin-6* (*Syt6*), a layer 6 marker, is expressed in the same layer as *Cyp26b1* in medial PFC at P8 (arrowhead). **N.** Schematic summary of the spatial expression patterns of *Cyp26b1* and *Aldh1a3* in medial PFC of early postnatal mouse brains. Scale bar, 1mm.

The tissue RA level is controlled both by its synthesis from vitamin A and by its degradation by CYP26 enzymes. Members of the aldehyde dehydrogenase 1 family are crucial for synthesizing RA, and two members of this family (ALDH1A2 and ALDH1A3) are expressed in early postnatal cortex; *Aldh1a2* is broadly expressed in the meninges^31^, whereas *Aldh1a3* is specifically expressed in the superficial layer of postnatal medial PFC as well as higher-order visual areas ^29,31^. We found that expression of *Aldh1a3* becomes detectable in medial PFC at P2 (Fig.1I). By P8, the expression was robust in layer 2 and extended more laterally (Fig.1J). A similar pattern of expression was found at P21 (Fig.1K) and P35 (Fig.1L). There also appears to be *Aldh1a3* expression in medial cortex in adult mice (Allen Brain Atlas). Outside of the frontal cortex, we did not detect *Cyp26b1* expression in more caudal parts of the neocortex including sensory areas at any developmental stage examined (Fig.S1D,E, S4D). *Cyp26b1* was also expressed in the piriform cortex, the amygdala, CA3 and the hilus regions of the hippocampus, and the globus pallidus (Fig.S1C-E). Expression in these regions started during embryogenesis (Fig.S1B,C) and continued into adulthood (Allen Brain Atlas). *Cyp26b1* was not detected in either medial (MGE) or lateral (LGE) ganglionic eminence (Fig.S1A).

In summary, in the medial part of early postnatal PFC, RA is produced by cells in the superficial layer, whereas CYP26B1, a RA degrading enzyme or an “RA sink”, is located in layer 6 (summarized in Fig.1N). These results suggest that RA signaling is spatially and temporally controlled, and that this regulation might play a role in development of medial PFC.

### Parvalbumin-expressing interneurons in medial PFC respond to retinoic acid during early postnatal development

The spatiotemporal expression pattern of *Cyp26b1* prompted us to explore the cellular targets of RA signaling during postnatal PFC development. In order to determine the populations of cells that respond to RA, we analyzed the *RARE-LacZ* transgene, an indicator of the transcriptional activity of the RA receptor, RAR (retinoic acid receptor)/ RXR (retinoid X receptor) heterodimers ^32^. At P0, expression of β-galactosidase (β-gal) was barely detectable in medial PFC except in radial glial fibers (Fig.2A). At P7, a small number of β-gal-positive cells were detected in medial PFC, mostly in layer 5 (Fig.2BE). These cells were also positive for SOX6, a marker for GABAergic interneurons derived from MGE^33^ (Fig.2B). This was still the case at P14, when the number of β-gal-positive cells greatly increased (Fig.2F-I); analysis of three transgenic brains revealed that 91% of β-gal-positive cells in layer 5 were also SOX6-positive (Fig.2N). Markers of other neuronal types, including SP8 (Fig.2C,J; caudal ganglionic eminence (CGE)-derived cortical interneurons)^34^, CTIP2 (Fig.2D,K; layer 5 subcerebral projection neurons as well as some interneurons)^35–37^ and TBR1 (Fig.2E,L; layer 6 corticothalamic projection neurons^38^) did not overlap with β-gal, indicating that these types of neurons do not express the molecular machinery necessary to respond to RA via RAR/RXR heterodimers. In summary, MGE-derived interneurons in layer 5 are the main responders to RA within early postnatal medial PFC.

**Fig.2.**
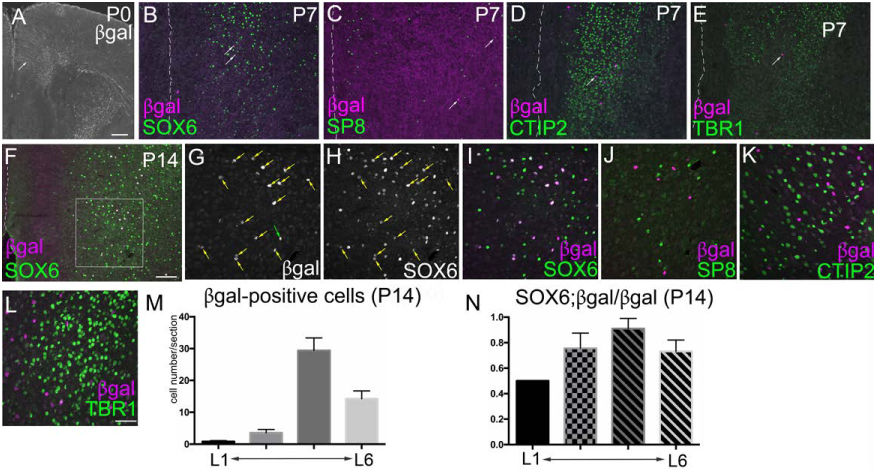
**RA signaling in early postnatal PFC** In all sections, the right hemisphere is shown, and the white dashed line marks the medial surface of the frontal cortex. Immunostaining for β-gal on frontal sections of P0 (A), P7 (B-E) and P14 (F-K) brains of *RARE-LacZ* transgenic mice. **A.** At P0, β-gal expression is only found in the radial glial fibers (arrow). **B-E.** At P7, a few β-gal-positive cells are found in the medial PFC, most of which are also SOX6-positive (B; white arrows show double-labeled cells). β-gal does not overlap with SP8 (C), CTIP2 (D) or TBR1 (E). **F-L.** At P14, the number of β-gal-positive cells dramatically increases from P7, and most of these cells are SOX6-positive. F is a 10x image including the medial surface of the brain, and G-I are 40x confocal images of the same region outlined in the square in F. I is the merged image of G and H. Yellow arrow in G,H show SOX6/β-gal double-positive cells, and green arrow in G shows a rare, β-gal-positive, SOX6-negative cell. (J-L) β-gal does not overlap with SP8, CTIP2 or TBR1. **M.** Average number of βgal-positive cells per section by layers (mean ± SEM). L1, layer 1 as marked by sparse labeling in DAPI staining. L6, layer 6 as marked by TBR1 staining. The two middle columns represent equal-width bins between layer 1 and layer 6, and roughly correspond to layers 2/3 and layer 4/5, respectively. Because most β-gal-positive cells are immediately above layer 6, and layer 4 is thin in medial PFC (Fig.S6A), the highest peak in the third column likely represents layer 5. **M**. The ratio of SOX6; β-gal-doublepositive cells among β-gal-positive cells are shown by layers (mean ± SEM). Scale bars, A: 200*μ*m, B-F:100*μ*m, G-L: 50*μ*m.

Most MGE-derived interneurons in the adult neocortex express either somatostatin (SST) or parvalbumin (PV)^39,40^. A majority of these interneurons complete their tangential migration into the neocortex by birth^41,42^. However, PV protein or *Pvalb* mRNA is not expressed until much later, indicating that the expression of PV or *Pvalb* is a useful marker of maturation for this lineage of cells. At P8, *Pvalb* mRNA was already abundant in lateral portion of the cortex including motor area (Fig.3A, double arrows), but not in medial PFC (Fig.3A arrow). Thus, maturation of PV neurons is delayed in medial PFC. By P14, cells expressing *Pvalb* mRNA or PV protein were clearly detectable in medial PFC, mainly in layer 5 (Fig.3B, arrow), which further increased by P21 (Fig.3C, arrow).

**Fig.3.**
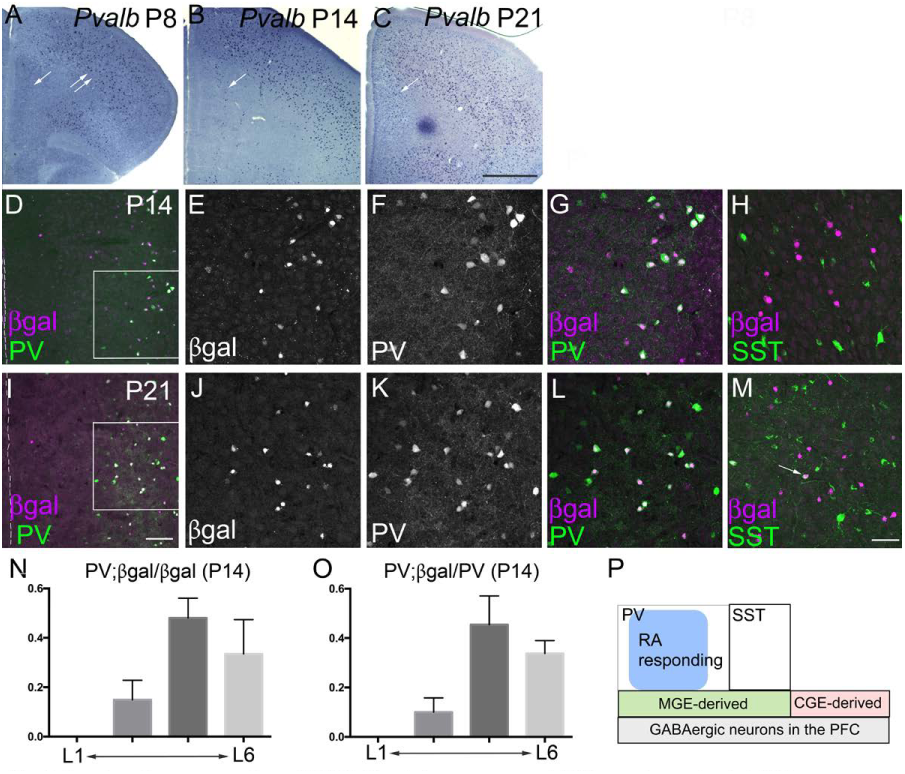
**Overlapping expression of *RARE-LacZ* transgene and PV in early postnatal PFC** **A-C.** In situ hybridization for *Pvalb* mRNA on frontal sections of control mice at P8 (A), P14 (B) and P21 (C). In medial PFC, *Pvalb* is undetectable at P8, but is robustly expressed at P14, which further increases by P21 (A-C, single arrow). **D-M.** Double immunostaining for β-gal and PV or β-gal and SST on frontal sections of P14 (D-H) and P21 (I-M) brains of *RARE-LacZ* transgenic mice. D is a 10x image including the medial surface of the brain, and E-G and J-L are 40x confocal images of the same region outlined in the square in D and I, respectively. G and L are merged images of E and F, and J and K, respectively. H and M are merged 40x confocal images of the medial PFC. Note the heavy overlap between β-gal and PV and little overlap between β-gal and SST (white arrow in M indicates a very rare double-positive cells). **N.** The ratios of PV; β-galdouble-positive cells among β-gal-positive cells in P14 medial PFC are shown by layers (mean ± SEM). **O.** The ratios of PV; β-gal-double-positive cells among PV-positive cells in P14 medial PFC are shown by layers (mean ± SEM). **P.** A schematic summary of the interneuron populations in medial PFC. Based on the results of this study, a subpopulation of PV interneurons respond to RA via RAR/RXR receptor complex. Scale bar, A-C: 1mm, D,I: 100μm, E-H, J-M: 50μm

In order to determine if RA-responding cells are restricted to either PV or SST neurons, we next analyzed *RARE-LacZ* transgenic mice to determine the co-localization of β-gal and PV (Fig.3D-G, I-L) as well as β-gal and SST (Fig.3H,M) at P14 and P21. We found a heavy overlap between PV and β-gal; in three P14 brains (Fig.3D-G), 48% percent of β-gal-positive cells in layer 5 were also PV-positive (Fig.3N). In turn, 45% of PV-positive cells in layer 5 were also β-gal-positive (Fig.3O). Similar patterns were found at P21 (Fig.3I-L). In a sharp contrast, SST show very little overlap with β-gal from P8 through P67 (Fig.3H,M and data not shown); the ratio of SST/β-gal double-positive cells to SST-positive cells was 2.9% (3 out of 103) at P14 and 5.0% (6 out of 121) at P21. In medial PFC, cells expressing SST protein or *Sst* mRNA appeared appeared between P0 and P4; by P7, ∼40% of MGE-derived interneurons that express SOX6 also expressed *Sst* mRNA, and this ratio stayed constant until P59 (Fig.S2). Thus, by P7, most SST neurons in medial PFC already express *Sst* mRNA, and that MGE-derived interneurons that are β-gal-positive and SST-negative at P14 or P21 (Fig.3H,M) are most likely to be PV neurons (summarized in Fig.3P).

### *Cyp26b1* is required for normal development of parvalbumin-expressing interneurons in medial PFC

Because of the strong correlation between the expression of PV and the cell’s responsiveness to RA, we hypothesized that development of PV interneurons in medial PFC is regulated by RA signaling and that this regulation depends on CYP26B1. To test this, we generated conditional *Cyp26b1* mutant mice in which *Cyp26b1* is deleted in the PFC. Because the expression of *Cyp26b1* is highly specific to layer 6 in the frontal cortex, we used *Synaptotagmin-6 Cre* (*Syt6-Cre*) driver mice^43,44^. Expression *Syt6* is specific to layer 6 in the neocortex (Fig.1M), and *Syt6-Cre* mice allow recombination in layer 6 corticothalamic projection neurons in the frontal cortex including the medial PFC (Allen Brain Atlas; http://connectivity.brain-map.org). To validate the usefulness of *Syt6-Cre* in knocking out *Cyp26b1* in layer 6 of the postnatal frontal cortex but not in other, potentially relevant *Cyp26b1*-expressing cell populations, we first bred the Cre mice with ZSGreen Cre reporter mice (Ai6)^45^. At E12.5, *Sy6-Cre*; *Ai6* brains showed ZSGR expression in the preplate and the meninges but not in other parts of the cortex or in ganglionic eminences (Fig.S3A). Cortical expression of ZSGreen started to be detected at E14.5 (Fig.S3B). At P0, robust signs of recombination were seen in layer 6 of the frontal cortex but not in more caudal cortex (Fig.S3C,D), amygdala (Fig.S3E) or CA3 and hilus regions of the hippocampus (Fig.S3D). Thus, we predicted that the *Syt6-Cre* mice would cause specific deletion of *Cyp26b1* in layer 6 cells of the frontal cortex including medial PFC. In *Syt6-Cre/+*; *Cyp26b1^flox/flox^* (*Cyp26b1 CKO*) mice, *Cyp26b1* mRNA was not detected in medial PFC (compare Fig.S4B and H as well as C and I), whereas expression in other brain regions including the hippocampus (Fig.S4D,J), agranular insula (Fig.S4C,I), globus pallidus (Fig.S4E,K) or amygdala (Fig.S4F,L) was not affected, confirming an efficient and specific deletion of *Cyp26b1*.

We then counted *Pvalb* mRNA-expressing neurons and compared the numbers between *Cyp26b1* CKO mice and their littermate wild-type controls at P14 and P21. At both stages, the density of *Pvalb*-positive cells was significantly increased in medial PFC of *Cyp26b1* mutant mice (Fig.4C; “total”, Fig.S5J; “total”). When the PFC was divided into superficial and the deep layers, the difference was seen only in the deeper half of the medial PFC, consistent with the distribution of β-gal expressing cells. With repeated measures 2-way ANOVA, we also detected significant difference both between layers and between genotypes (see legend in Fig.4). In the motor cortex, we detected significant increase in the density of *Pvalb*-expressing cells in deep layers of *Cyp26b1* CKO compared with controls by using paired t-tests (Fig.4D), but the difference was not significant with repeated measures 2-way ANOVA. More laterally, the putative somatosensory area also did not show significant difference in density (Fig.4E), suggesting that in the CKO mice, the residual *Cyp26b1* expression in layer 5 of lateral frontal cortex (Fig.S4I) might have spared the normal density of *Pvalb*-expressing cells in lateral cortex. In the medial PFC, normal expression of *Cyp26b1* is limited to layer 6, and CKO mice had no residual expression in more superficial layers. In contrast to *Pvalb*-expressing neurons, the density of *Sst*-, *Vip*- (derived from CGE) or *Lhx6*- (a general marker for MGE-derived interneurons) expressing neurons was not significantly different in the CKO mice compared with controls (Fig.S5A-I,K). Furthermore, in adult brains, *Pvalb*-, *Sst*-, and *Vip*-expressing cells did not show significant difference between the CKO and control mice (Fig.S5L-M). Together, these results indicate that transient expression of *Cyp26b1* in layer 6 of medial PFC is specifically required for controlling the development of PV interneurons. This is consistent with disproportionately high percentage of PV neurons responding to RA compared with SST neurons (Fig.3). The lack of significant change in the density of *Sst-*, *Vip*- or *Lhx6-*expressing cells suggests that CYP26B1 does not control the fate specification or the number of each interneuron type, but rather the rate of maturation of PV lineage cells specifically.

**Fig.4.**
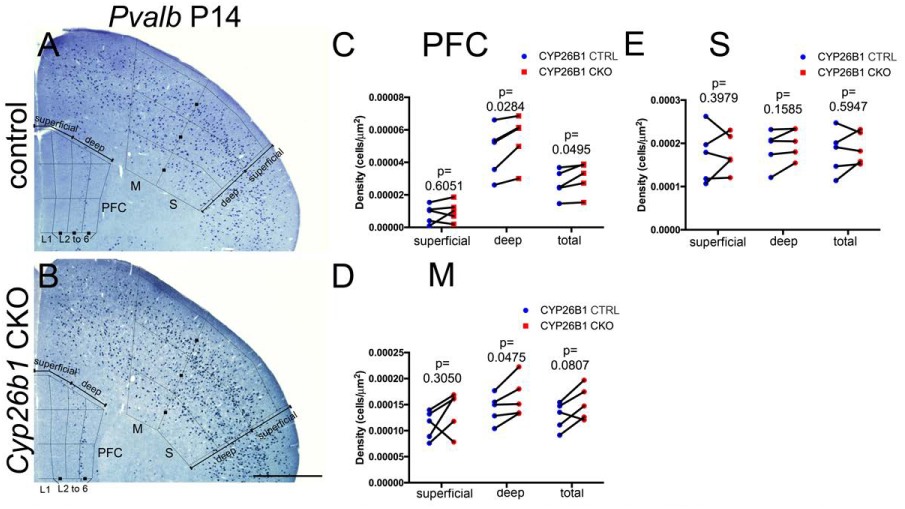
**Increased *Pvalb*-expressing interneurons in medial PFC of *Cyp26b1* knockout mice** **A,B.** In situ hybridization of frontal sections of P14 *Cyp26b1* conditional knockout mice (**B**) and littermate controls (**A**). Expression of *Pvalb* mRNA is shown. See Methods on binning of the medial PFC. Numbers of *Pvalb*-positive cells in the two superficial bins and two deep bins were added together and compared separately between *Cyp26b1* mutants and littermate controls. **C**-**E** show the result of paired ratio t-tests for cell counts in the medial PFC, motor cortex (D) and somatosensory cortex (E), all on the same frontal sections. Each line connecting red and blue dots represents a pair of brains analyzed in the same experiment (n=5). p-values of ratio paired t-tests for each layer (superficial, deep, total) are shown. In repeated measures two-way ANOVA, the p-values for layer (superficial vs deep), pair (between control and knockout), and interactions (between layer and pair) are 0.0017, 0.0325 and 0.1416 (P14 in PFC); 0.0441, 0.0961 and 0.7807 (P14 in motor cortex), 0.1771, 0.4751 and 0.5496 (P14 in somatosensory cortex), respectively. Scale bar, 1mm. L1: layer 1

### *Cyp26b1* is not expressed in early postnatal medial PFC in the absence of thalamus-PFC connectivity

Because *Cyp26b1* starts to be expressed in medial and ventral PFC when the reciprocal connections between the thalamus and the cortex are being established, we next asked if the normal expression of *Cyp26b1* depends on this connectivity. In our previous study, thalamus-specific deletion of the homeobox gene *Gbx2* resulted in severe deficiency of thalamocortical and corticothalamic projections in sensory areas^46^. Similar to sensory cortex, the PFC also showed a significant reduction in the staining of NetrinG1, a marker of thalamocortical axons^46,47^ in *Gbx2* mutant mice at E16.5 (Fig.S6A,B). Placement of DiI crystals into the medial PFC of *Gbx2* mutants and wild type mice at P14 revealed that both retrograde labeling of thalamic neurons and anterograde labeling of corticothalamic axons were severely attenuated in *Gbx2* mutants (Fig.S6C-F). These results demonstrate a robust reduction of reciprocal connectivity between the thalamus and PFC in *Gbx2* mutant mice.

We then tested if expression of *Cyp26b1* in the PFC is altered in *Gbx2* mutant mice. Already at P2, the mutant cortex lacked the expression of *Cyp26b1* in medial and ventral PFC (Fig.5A,F), demonstrating that the onset of *Cyp26b1* expression requires thalamocortical interactions. The deficiency of *Cyp26b1* expression continued until P21, when the medial PFC expression of *Cyp26b1* normally started to decline (Fig.5D,I). In contrast, expression of *Cyp26b1* showed no alterations in layer 5 of the lateral frontal cortex including the motor areas and agranular insula (Fig.5A-J). The layer 6 marker, *Syt6* was still highly expressed in the frontal cortex of *Gbx2* mutants (Fig.5K,N), making it unlikely that cell loss in layer 6 caused the reduced expression of *Cyp26b1* in *Gbx2* mutants. Lastly, *Aldh1a3,* which normally shows a similar onset of expression in medial PFC to that of *Cyp26b1*, was qualitatively unaffected in *Gbx2* mutants (Fig.5L,O). In summary, transient expression of *Cyp26b1* in layer 6 of medial PFC was dependent on the connections between the thalamus and the cortex (summarized in Fig.5M,P). Putting together the role of *Cyp26b1* in development of PV neurons in medial PFC, our results collectively demonstrate a novel, indirect role of the thalamus in regulating the neocortical interneuron development.

**Fig.5.**
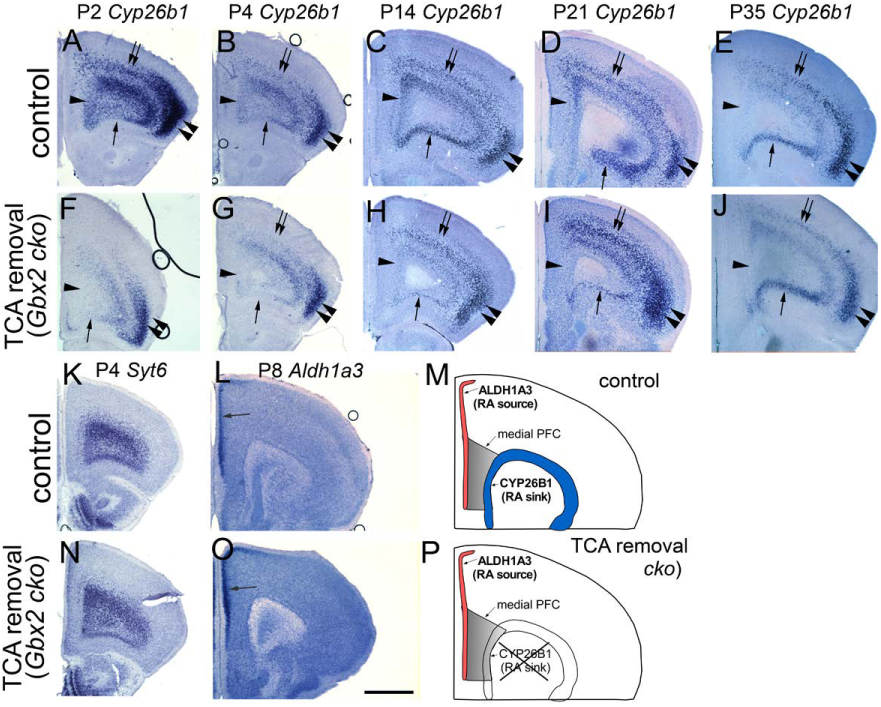
**Transient expression of *Cyp26b1* in the PFC does not occur in the absence of thalamus-cortex interactions in *Gbx2* mutant mice.** **A-J.** in situ hybridization of frontal sections through PFC at various postnatal stages with a *Cyp26b1* probe. In control mice (A-E), *Cyp26b1* expression starts at P2 in medial (A, arrowhead) and ventral (A, single arrow) PFC, and continues until P14 (C). At P21, expression in medial PFC is reduced (D), and is no longer detectable at P35 (E). In addition to medial and ventral PFC, *Cyp26b1* is also expressed in lateral frontal cortex including the motor and somatosensory areas (A-E, double arrows) and agranular insula (A-E, double arrowheads). In *Gbx2* mutant mice (F-J), expression of *Cyp26b1* is not induced in medial or ventral PFC at P2 as well as at later stages, although ventral PFC does not appear to be affected at P14 and later. Expression in more superficial layer of lateral cortex (G-J, double arrows and double arrowheads) is not affected in *Gbx2* mutant mice. **K,N.** Expression of the layer 6 marker *Syt6* is not affected in *Gbx2* mutant mice. **L,O.** Expression of *Aldh1a3* in layer 2 of medial PFC and anterior cingulate cortex (arrow) is not affected in *Gbx2* mutant mice. scale bar, 1mm. **M,P.** Summary schematic for this figure

### Lack of thalamus-PFC connectivity results in early aberrancy of radial positioning of MGE-derived interneurons

In sensory cortex, thalamocortical afferents affect the development of interneurons by a variety of mechanisms^12–15^. However, roles of the thalamus in the development of PFC interneurons are unknown. Hence, we analyzed the distribution of MGE-derived interneurons in medial PFC both at P21 and P0. At P21, densities of *Pvalb*- and *Sst*- expressing interneurons in medial PFC, specifically in the middle layers and not in the most superficial and the deepest layers, were significantly reduced in *Gbx2* mutant mice (Fig.6A-C). Already at P0, LHX6-expressing, MGE-derived interneurons showed an altered radial distribution in medial PFC (Fig.6D-G); there was an increase in the density of LHX6-positive cells in layer 6 and below, and a decrease in the middle layers (Fig.6F), while the total density was not significantly different between the two genotypes (Fig.6G). These results on the neonatal PFC are remarkably similar to a recent report on sensory and motor cortex of thalamus-specific *Gbx2* mutant mice^12^. Thus, there is an early, cortex-wide role of thalamocortical projections in controlling the radial distribution of MGE-derived interneurons. Because the altered neuronal positioning had already occurred at P0 in *Gbx2* mutant mice, which is before the onset of *Cyp26b1* expression in medial PFC, these early roles are likely to be independent of the later role of the thalamus in regulating the development of PV interneurons via *Cyp26b1*.

**Fig.6.**
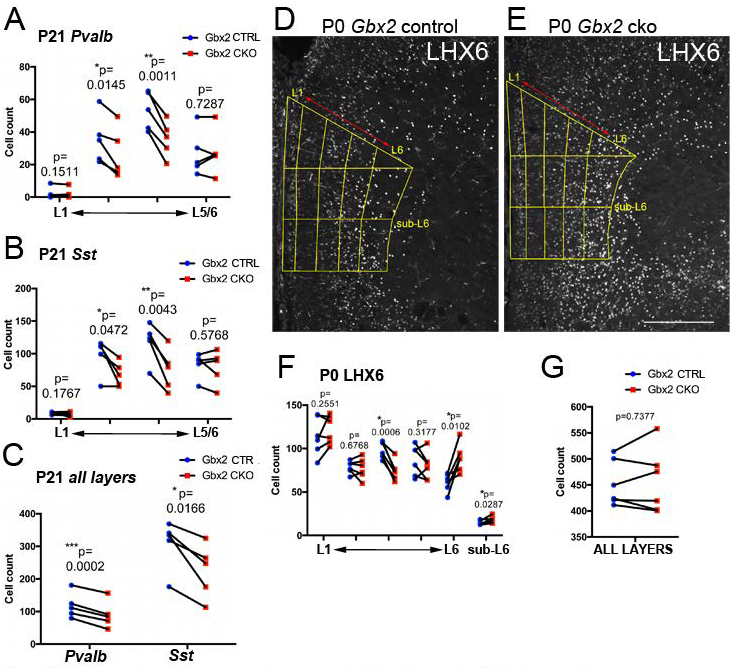
**Abnormal numbers and radial distribution of MGE-derived interneurons in the medial PFC of P21 and neonatal *Gbx2* mutant mice** **A-C.** Comparison of *Pvalb*-positive and *Sst*-positive cells in *Gbx2* mutants (red dots) and wild-type littermates (blue dots) in medial PFC at P21. Each line connecting red and blue dots represents a pair of brains analyzed in the same experiment (n=5). A and B show comparison of laminar distribution of *Pvalb*-expressing and *Sst*-expressing neurons, respectively, in *Gbx2* mutant mice and littermate controls. Layer 1 was defined as the cell-sparse layer detected by DAPI staining. The remaining cortical wall was equally divided into three layers. The deepest layer (shown as “L5/6”) contains the entire layer 6 and the deep part of layer 5. C shows the total number of *Pvalb*- and *Sst*expressing neurons in all layers. P-values of paired t-test for individual layer are shown. In repeated measures two-way ANOVA, the p-values for layer, pair (between control and knockout), and interactions (between layer and pair) are <0.0001, 0.0002 and 0.0001 (*Pvalb*); <0.0001, 0.0247 and 0.0001 (*Sst*), respectively. **D-E.** Representative images of immunostaining for LHX6 in medial PFC of wild-type (D) and *Gbx2* mutant (E) mice at P0. Binning is shown in yellow. Layer 1 (L1) was defined as the cell-sparse layer detected by DAPI staining. Layer 6 (L6) was defined as the layer with TBR1 staining on the same sections (not shown). The intervening region was equally divided into three layers. Sub-layer 6 was defined as the layer below layer 6. Scale bar, 200μm **F-G.** Comparison of LHX6-positive cells in *Gbx2* mutants (red dots) and wild-type littermates (blue dots) in medial PFC at P0. Each line connecting red and blue dots represents a pair of brains analyzed in the same experiment (n=5). **F** shows laminar distribution pattern. P-values of paired t-test for individual layer are shown. In repeated measures two-way ANOVA, the p-values for layer, pair (between control and knockout), and interactions (between layer and pair) are 0.0001, 0.5950 and 0.0021, respectively. **G** shows the total number of *Pvalb*- and *Sst*-expressing neurons in all layers. P-values of paired t-test are shown. “Sub-L6” was defined as the region below the expression domain of TBR1, which was stained in all immunostaining slides for a reference. *: p<0.05, **: p<0.005, ***: p<0.0005

### Induction of *Cyp26b1* by the thalamus is independent of transmitter release from thalamocortical projection neurons

How does the thalamus control the expression of *Cyp26b1* in layer 6 neurons in medial PFC at early postnatal stages? One likely cue that mediates the role of the thalamus is the transmitter release from thalamocortical axons. To test if the lack of transmitter release phenocopies the lack of the axon projections, we generated mutant mice in which tetanus toxin light chain (TeNT) is ectopically expressed specifically in thalamocortical projection neurons (Fig.7). At E16.5, expression of VAMP2, the cleavage target of TeNT, was dramatically reduced in thalamocortical axons expressing TeNT, while it was retained in corticofugal axons (Fig.7A-D, G-J). At P8, expression of *RORβ* in layer 4 of the primary somatosensory area was altered in TeNT-expressing mice, lacking the characteristic barrel-like pattern (Fig.7E,F). This is consistent with a recent study on *vGluT* mutants^48^ and indicates the role of transmitter release in the formation of normal cytoarchitecture of the primary sensory cortex. In the PFC, however, induction of *Cyp26b1* in the medial and ventral PFC was not qualitatively affected in TeNT-expressing mice (Fig.7K,L), implying a unique cellular mechanism that underlies the induction of *Cyp26b1* expression in early postnatal PFC.

**Fig.7.**
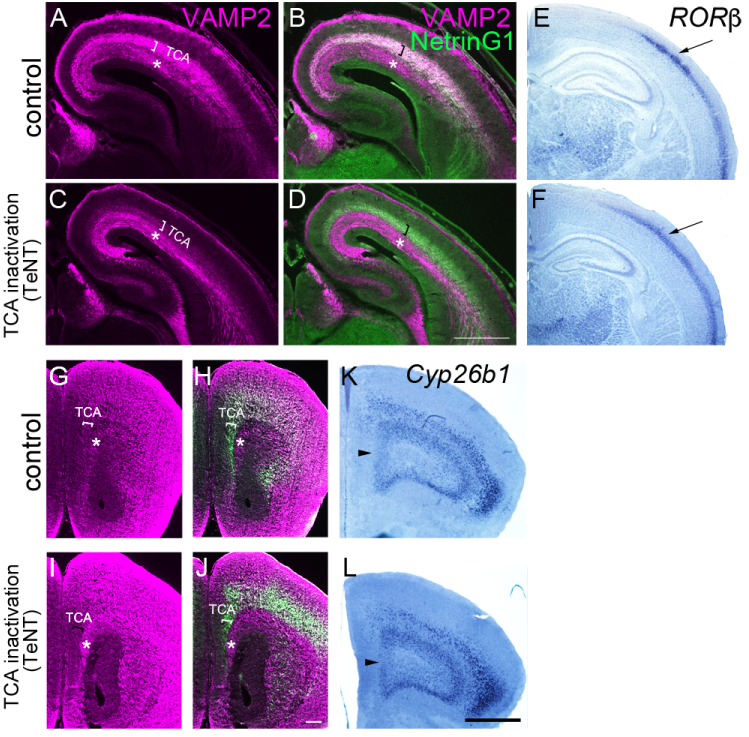
**Normal induction of *Cyp26b1* in PFC in mice expressing tetanus toxin light chain in thalamocortical axons** **A-D.** Immunostaining for VAMP2 on frontal sections of somatosensory cortex at E16.5 control (A,B) and mutant mice with ectopic expression of tetanus toxin light chain (TeNT) in thalamic neurons (C,D). TeNT expression leads to deletion of VAMP2 specifically in thalamocortical axons at E16.5. Thalamocortical axons are shown by NetrinG1 staining (B,D, green). In control brains, both thalamocortical (shown in asterisk in A-D) and corticofugal (shown in bracket in A-D) axons express VAMP2, whereas in TeNT-expressing mice, VAMP2 staining is specifically diminished in thalamocortical axons (C,D, asterisk). Scale bar, 500μm. **E,F.** Deletion of VAMP2 in thalamocortical axons results in the lack of the characteristic pattern of *RORβ* expression in the barrel field of primary somatosensory cortex at P8 (arrow), similar to the defect found in *Gbx2* mutant mice (Vue et al., 2013). **G-J.** Immunostaining for VAMP2 on frontal sections of prefrontal cortex at P0 control (G,H) and mutant mice with ectopic expression of tetanus toxin light chain (TeNT) in thalamic neurons (I,J). Similar to the somatosensory cortex, VAMP2 staining in thalamocortical axons is diminished in TeNT-expressing mice (bracket in I, J). Scale bar, 200μm. **K,L.** Expression of *Cyp26b1* in medial (arrowhead) and ventral PFC is intact in TeNT-expressing mice (L), similar to control (K), at P8. scale bar, 200μm (G,H,I,J) 1mm (E,F,K,L)

## Discussion

In this study, we first demonstrated that *Cyp26b1*, which encodes an RA-degrading enzyme and a critical regulator of retinoid signaling^49,50^, is expressed in developing PFC in a temporally and regionally specific manner. We also showed that PV-expressing interneurons are the major cell population that normally responds to RA via the RAR/RXR receptors. Conditional deletion of *Cyp26b1* in layer 6 of the frontal cortex resulted in an increased density of *Pvalb*-expressing neurons in deep layers of medial PFC during postnatal development. Expression of *Cyp26b1* in the PFC depended on the connections between the cortex and the thalamus, but not on the transmitter release from thalamocortical axons. These results demonstrate a unique regulatory role of the thalamus in postnatal development of PV interneurons in the PFC (summarized in Fig.8D-G).

**Fig.8.**
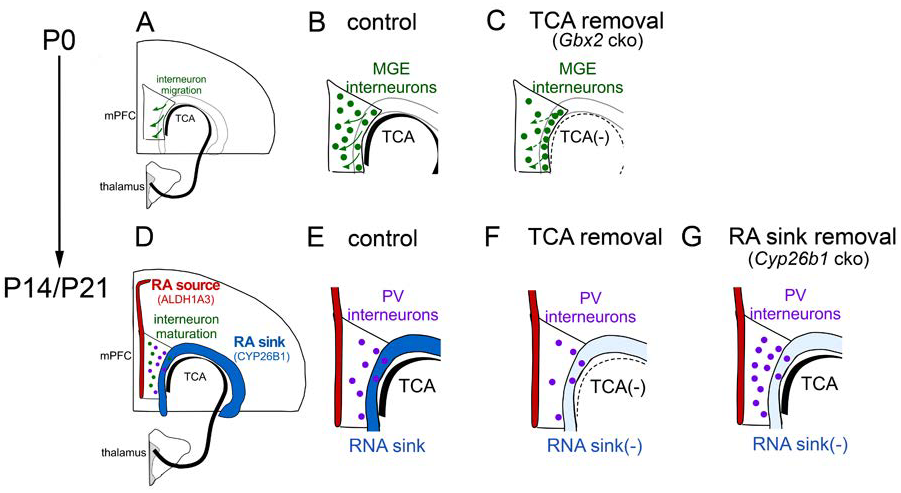
**Schematic diagrams of the current finding** **A-C.** Embryonic roles of thalamocortical axons as observed in neonatal mice. (A) Thalamocortical axons reach the medial PFC (mPFC) by E16.5 and control migration of medial ganglionic eminence (MGE)-derived interneurons. (B) In normal neonatal mice, MGE-derived interneurons have largely completed tangential migration to the mPFC and have taken proper laminar positioning by radial dispersion (arrows). (C) In thalamus-specific *Gbx2* mutant mice, radial positioning of MGE-derived interneurons are aberrant, resulting in their accumulation in layer 6 and below. **D-G.** Postnatal roles of thalamus-PFC interactions and retinoic acid (RA)-degrading enzyme CYP26B1 in the development of PV interneurons in the mPFC. (D) Early postnatal mPFC is positioned between the source of RA synthesis (layer 2, by ALDH1A3) and the RA-degrading “sink” (layer 6, by CYP26B1). Expression of both enzymes is induced early postnatally, but only *Cyp26b1* is dependent on the connections with the thalamus. The main cell population that responds to RA in postnatal mPFC are parvalbumin (PV) interneurons, and their development is controlled by CYP26B1. (E) In normal postnatal mice, PV neurons mature and start to express *Pvalb* mRNA and PV protein mainly in deep layers of mPFC between P7 and P14. (F) In thalamus-specific *Gbx2* mutant mice, *Cyp26b1* is not induced in mPFC. The number of both *Pvalb* and *Somatostatin* (*Sst*)-expressing neurons is reduced in the middle layers at least partly due to the earlier defects in radial dispersion (described in C). (G) In frontal cortex-specific *Cyp26b1* mutant mice, lack of the RA sink in mPFC leads to an increased number of neurons that express *Pvalb* mRNA or PV protein in deep layers.

### Roles of RA signaling in postnatal development of the medial PFC

In early embryonic brain, RA controls rostrocaudal patterning of the hindbrain and spinal cord by regulating the expression of *Hox* genes^21,22^. Cells in the subventricular zone of the embryonic LGE express the RA-synthesizing enzyme ALDH1A3, and *Aldh1a3* deficient mice had reduced expression of dopamine receptor D2 (*Drd2*) in nucleus accumbens^23^ and reduced expression of *Gad1* in embryonic GABAergic neurons in the striatum and the cortex^24^, demonstrating a crucial role of RA in early differentiation of embryonic GABAergic neurons. However, much less is known about the roles of RA signaling in postnatal brain development. Systemic administration of RA into early postnatal mice caused behavioral hyperactivity at the adult stage with an increased number of calbindin-expressing neurons in the cingulate cortex^51^. Thus, it was expected that RA has a role in early postnatal brain development. However, the role of RA in PFC development and the underlying cellular and molecular mechanisms had remained unknown.

Our current study revealed that in postnatal PFC, a significant subpopulation of PV interneurons are responsive to RA via RAR/RXR receptors, and that the lack of the RA-degrading enzyme, CYP26B1, causes an increase in the density of *Pvalb*expressing cells in medial PFC at P14 and P21. This phenotype should be independent of the proposed, earlier roles of RA in tangential migration of GABA neurons^52^, because 1) most MGE-derived interneurons have completed the tangential migration to the cortex by birth, 2) *Cyp26b1* is expressed only postnatally in the medial PFC and is not expressed in embryonic MGE or LGE (Fig.S1A,B), 3) *Syt6-Cre* mice do not cause recombination in MGE-derived interneurons (Fig.S3), and 4) in *Syt6^Cre/+^; Cyp26b1^flox/+^* mice, expression of *Cyp26b1* is not affected in regions outside of the frontal cortex, including the ventral telencephalon (Fig.S4).

In addition, the increased number of *Pvalb*-positive neurons in deep layers of medial PFC was not accompanied by their decrease in upper layers. This suggests that the radial dispersion of PV neurons, which follows their tangential migration, was also unaffected in *Cyp26b1* mutants.

At the adult stage, *Cyp26b1* mutants no longer showed a significant increase in the number *Pvalb*-positive neurons in medial PFC (Fig.S5L). In addition, the total number of MGE-derived interneurons marked by LHX6 did not show a significant change throughout postnatal development. Thus, the most likely role of *Cyp26b1* in postnatal development of PV neurons in the medial PFC is to slow their rate of maturation by suppressing RA signaling. It is known that neurotrophin (e.g., BDNF) signaling and neuronal activity, likely mediated by NMDA receptors, play a role in postnatal development of PV neurons^53–57^. Further studies are needed to investigate the relationship between these pathways and RA signaling. It will also be informative to analyze various aspects of maturation of PV neurons including intrinsic electrophysiological properties, formation of perineuronal nets and changes in gene expression^16,58^.

### Universal and area-specific roles of the thalamus in neocortical development

Previous studies have indicated that thalamocortical afferents have instructive roles in establishing area‐ and layer-specific gene expression as well as in promoting morphological differentiation of excitatory neurons in primary visual and somatosensory cortex^46,48,59,60^. Although the underlying cellular and molecular mechanisms have just begun to be elucidated, at least some of the effects appear to be dependent on the release of neurotransmitters from thalamocortical axons^48^. In addition to excitatory neurons, both SST‐ and PV‐ interneurons also depend on thalamic afferents for their maturation, likely via glutamatergic synaptic transmission^13–15^. In somatosensory, visual and motor areas, thalamic afferents also control radial positioning of MGE-derived interneurons before birth, in which the thalamus regulates the expression of the KCC2 co-transporter via the release of glutamate from thalamic axon terminals^12^. Are these roles universal throughout the cortex or is there area-specific regulation of cortical development by the thalamus? The thalamus is composed of dozens of nuclei with distinct gene expression profiles^19,61^, and each thalamic nucleus has a unique patterns of axonal projections to different cortical areas^20^. The neocortex is also regionally specified before the arrival of the thalamic axons^62–64^, and exhibits a rostral-to-caudal and lateral-to-medial neurogenic gradients^65^. Thus, it is expected that the nature of interactions with the thalamus varies between different cortical areas.

In fact, expression of ROR*β* and *Lmo4*, which showed abnormal patterns in primary sensory cortex of thalamus-specific *Gbx2* mutant mice, was not impaired in the medial PFC (Fig.S7). Instead, our current study has revealed a frontal cortex-specific regulation of *Cyp26b1* by the thalamus at early postnatal stages; loss of axons between the thalamus and the cortex in *Gbx2* mutants prevented *Cyp26B1* from being expressed in layer 6 of the frontal cortex. Interestingly, our TeNT model, in which the vesicle release from TCAs was blocked, was insufficient for recreating the loss of *Cyp26B1* seen in the morphological deficiency of thalamocortical connectivity. Therefore, the role of the thalamus in inducing *Cyp26b1* expression in layer 6 PFC neurons is likely independent of transmitter release that involves VAMP2 functions.

The current study also revealed an early role of the thalamus in regulating the radial positioning of MGE-derived interneurons in neonatal PFC. The altered positioning of interneurons was highly reminiscent of what was found in sensory and motor areas^12^. This indicates that the early role of the thalamus in controlling the radial positions of cortical interneurons is shared between many neocortical areas (summarized in Fig.8AC). Importantly, radial positioning of cortical interneurons is compromised also in *Lhx6* mutant mice, which suggests that the tangential-to-radial switch of interneuron migration in the cortex requires coordination of intrinsic and extrinsic signaling mechanisms during embryonic development.

It is intriguing that in postnatal *Gbx2* mutant mice, even in the absence of normal induction of *Cyp26b1* in medial PFC, the density of *Pvalb*-expressing neurons was decreased, not increased, unlike in *Cyp26b1* mutants. This suggests that the thalamus plays additional roles in postnatal development of PV neurons other than by inducing *Cyp26b1*, which include the support of their survival. Thus, it will be important to determine if other known roles of the thalamus in postnatal regulation of the development of PV neurons in sensory cortex ^13–15^ also apply to the PFC.

### Functional implications of altered PV neuron development in *Cyp26b1* mutant mice

At the systems level, RA regulates cortical synchrony during sleep^66^, memory and cognitive behaviors^67–69^. In addition, aberrant RA signaling is associated with multiple psychiatric disorders including schizophrenia, bipolar disorder and depression in humans^70–73^. Because many psychiatric disorders are predisposed by aberrant brain development, understanding how RA functions in early postnatal brain development is important for determining the long-term consequences of the perturbations of this signaling pathway. We found that in the absence of *Cyp26b1* in frontal cortex during a discrete period of postnatal development, the medial PFC had more *Pvalb*-expressing interneurons than in control mice. It is well established that PV neurons orchestrate activity in local circuits, which leads to oscillatory synchronous network activity in the gamma-band^74,75^. Synchronous gamma-band activity in medial PFC is associated with the successful operation of working memory. In a mouse genetic model of schizophrenia that replicates the human the 22q11.2 microdeletion syndrome^76^, gamma synchrony and working memory performance were impaired^77^, so was the development of PV interneurons^3,78–80^. Furthermore, mutations in the *Disc1* gene, a significant risk factor for schizophrenia and other disorders including depression and bipolar disorder in humans, cause alterations of PV neurons and reduction of gamma oscillations in mice^81^. These findings link PV interneuron abnormalities to changes in prefrontal synchrony and working memory impairment in mouse models of neuropsychiatric disorders. Mutation of CYP26B1 in humans is a risk factor for schizophrenia that reaches genome wide significance^82,83^. Therefore, it will be interesting to test whether the aberrant time-course of PV neuron development in medial PFC causes altered synchrony and impairment of cognitive behaviors in *Cyp26b1* mutant mice.

## Materials and methods

### Mice

*RARE-LacZ* transgenic mice^32^ were obtained from Jackson Laboratory (stock number: 008477) and were kept in the CD1 background. Frontal cortex-specific *Cyp26b1* mutant mice were generated using BAC (bacterial artificial chromosome) *Syt6-Cre* mice (GENSAT)^43,44^. Although the endogenous *Syt6* gene is expressed ubiquitously in layer 6 of the neocortex, the BAC Cre line caused recombination specifically in the frontal cortex. We crossed *Syt6^Cre/+^; Cyp26b1^flox/+^* mice and *Cyp26b1^flox/flox^* mice to generate the conditional mutants. *Cyp26b1^flox/flox^* mice^84^ were developed by Dr. Hiroshi Hamada’s laboratory (RIKEN CDB, Kobe, Japan) and obtained from Dr. Maria Morasso (NIAMS, Bethesda, MD). *Rosa26^stop-ZSGreen/+^* (*Ai6*) mice^45^ were obtained from Jackson Laboratory (stock number: 007906). Thalamus-specific *Gbx2* mutant mice were generated by crossing *Olig3^Cre/+^; Gbx2^null/+^* mice and *Gbx2^flox/flox^* mice as described^46^. *Gbx2^flox/flox^* mice were obtained from Jackson Laboratory^85^. *Olig3^Cre/+^* mice were described previously^86,87^. The *Gbx2^null^* allele was generated by crossing *Gbx2^flox/flox^* mice with the *CMV-Cre* germ-line deleter mice (Jackson Laboratory, stock number: 003465). Mice that express tetanus toxin light chain in thalamic neurons were generated by crossing *Olig3^Cre/+^* mice and *Rosa26^stop-TeNT/stop-TeNT^* mice^88^.

### In situ hybridization

cDNAs for the following genes were used: *Cyp26b1*, *Aldh1a3*, *Syt6, Lmo4* (Open Biosystems), *Pvalb*, *Sst*, *Vip* (obtained from Dr. Rob Machold*), Lhx6* (obtained from Dr. John Rubenstein), *RORβ*, (obtained from Dr. Michael Becker-Andre). Postnatal pups were perfused with 4% paraformaldehyde/0.1M phosphate buffer and the heads were post-fixed until needed. Brains were then taken out of the skull, washed in 0.1M phosphate buffer for 20min and were sunk in 30% sucrose/0.1M phosphate buffer. Coronal sections were cut with a sliding microtome at 50μm-thickness (Leica) or with a cryostat at 20μm‐ (P2 or younger) or 40μm-thickness (P4 or older) and were mounted on glass slides (Super Frost Plus, Fisher). In situ hybridization was carried out as described^89^.

### Immunohistochemistry

Brains were taken out immediately after perfusion and were postfixed for 1 hour (P0), 1-2 hours (P4-P14) or 1-4 hours (P21). After the post-fixation, the brains were washed in 0.1M phosphate buffer for 20min and were sunk in 30% sucrose/0.1M phosphate buffer. Sections were cut in the same way as those used for in situ hybridization. The following primary antibodies were used: β-galactosidase (Cappel, 55976, goat, 1:100; Abcam, chicken, 1:500), SOX6 (Abcam, rabbit, 1:100), SP8 (Santa Cruz, sc-104661, goat, 1:100), CTIP2 (Abcam ab18465, rat, 1:200), TBR1 (Abcam, ab31940, rabbit, 1:200; Millipore, AB2261, chicken, 1:200), PV (SWANT, rabbit, 1:500), SST (Millipore, MAB354, rat, 1:100), LHX6 (Santa Cruz, sc-271433, mouse, 1:50), VAMP2 (Synaptic Systems, 104 202, rabbit, 1:200), NetrinG1 (R&D Systems, goat, 1:100). Secondary antibodies conjugated with Cy2, Cy3 or Cy5 were obtained from Jackson ImmunoResearch.

### Combined in situ hybridization (*Sst*) and immunohistochemistry (SOX6)

In situ hybridization was carried out as described above, except that prior to proteinase K treatment, sections were incubated with 0.3% hydrogen peroxide/PBS for 10 minutes to block endogenous peroxidase activity. After the hybridization, slides were washed as described above, and were incubated for 30 minutes with anti-DIG antibody conjugated with peroxidase, followed by two washes in PBS and one wash with TNT solution (TNT: 0.1M Tris pH7.5, 0.15M NaCl, 0.05% Tween 20). Sections were then incubated for ∼30 min in Tyramide Plus fluorescein (ThermoFisher) diluted at 1:250, washed twice in TNT solution and post-fixed. Thereafter, immunostaining with anti-SOX6 antibody was carried out as described above.

### Imaging/Binning

For cell counting, images of sections that underwent in situ hybridization or immunostaining were taken using an upright microscope (Nikon E800) with a 2x (in situ hybridization) or a 4x (immunostaining) objective using a digital CCD camera (Retiga EXi, QImaging) and the OpenLab software. The range of coronal sections to be included in the analysis of medial PFC was defined rostrally by the presence of the lateral ventricle and caudally by the presence of the anterior forceps of the corpus callosum. Once the sections were selected, their images were scrambled and each file was given a randomized number so that the person who performed the subsequent analysis was made blind to the genotypes. Cell counting was performed in the putative prelimbic (PL) and infralimbic (IL) areas of the medial PFC (Allen Brain Atlas; ^90,91^). Images were stacked using Adobe Photoshop and subsequently binned to distinguish the parts of the medial PFC as shown in Fig.4; each bin is 500μm-high at the medial surface. For sections with in situ hybridization, the most superficial layer (layer 1) was defined as the cell-sparse layer on DAPI staining. The remaining cortical wall was divided into three bins with equal widths, resulting in 4 layers of bins. Layer 1 and the layer underneath it were grouped together and were named superficial layers and the remaining two layers were named the deep layers. For immunostaining, we used anti-TBR1 antibody for all slides in Cy5 channel and used it as a reference marker of layer 6. Then, the areas excluding layers 1 and 6 were equally divided into three parts, resulting in five layers of bins (shown in Fig.6). For Fig.2 and Fig.3, the portion between layer 1 and layer 6 was divided into two equal parts, resulting in four layers of bins. For high-magnification images shown in Fig.2 and Fig.3, an Olympus FluoView 1000 confocal microscope was used (40x oil, NA=1.25).

### Cell Counting

In imageJ, the Image-based Tool for Counting Nuclei (ITCN) plugin, from the Center for Bio-Image Informatics, UC Santa Barbara, was used to count cells and measure the area of each bin.

### Statistical analysis

Paired ratio t-test was used for comparing cell counts between *Gbx2* cko mice and wild-type littermates as well as *Cyp26b1* cko and wild-type littermates. For data shown in Fig.4, Fig.S5 and Fig.6, we also performed repeated measures 2-way ANOVA using Prism (versions 6 and 7, GraphPad). Comparison was made between two brains processed and analyzed in the same experiment. Graphs were generated using Prism.

### Axon tracing

Small crystals of 1,1′-dioctadecyl-3,3,3′3′-tetramethylindocarbocyanine perchlorate (DiI) were placed on the medial surface of frontal cortex of PFA-fixed P14 *Gbx2* conditional mutant brains and their control littermates. After incubation of the brains in PFA at 37℃ for 2 weeks, we cut sections at 150μm with a vibrating microtome (Vibratome), counter-stained the sections and mounted them on glass slides for imaging.

## Acknowledgments

We also thank Steven McLoon, Goichi Miyoshi and Timothy Monko for helpful comments and discussions, and Thomas Bao, Samantha Dabruzzi, Shaylene McCue, Morgan McCullough and Carmen Tso for technical assistance, and University of Minnesota Vision Core and Paulo Kofuji for allowing us to use their Olympus confocal microscope. Zachery Werkhoven and Melody Lee contributed to the initial phase of the project. We thank Martyn Goulding for providing *Rosa26^stop-TeNT/+^* mice, Hiroshi Hamada (RIKEN Center for Developmental Biology) and Maria Morasso (NIH) for *Cyp26b1*^flox/+^ mice and Eric Turner (University of Washington) for *Syt6*^Cre/+^ mice, Rob MacHold for *Pvalb, Vip* and *Sst* cDNAs, John Rubenstein for *Lhx6* cDNA. This work was supported by grants to Y.N. from NIH (R21MH105759), Brain and Behavior Research Foundation (Essel Investigator), Winston and Maxim Wallin Neuroscience Discovery Fund and Academic Health Center of the University of Minnesota.

## LEGENDS TO SUPPLEMENTAL FIGURES

**Fig.S1 Normal expression pattern of *Cyp26b1* in the mouse forebrain**

In situ hybridization on coronal sections is shown. **A,B.** Expression is not detected in LGE or MGE at E14.5, but is already found in the hippocampus (A,B, arrowhead), septum (A, arrow), globus pallidus (“GP” in B) and amygdala (“Amy” in B). B is more caudal than A. **C.** At E16.5 (C), *Cyp26b1* is detected in hippocampus (arrowhead), piriform cortex (“Pir”), globus pallidus (“GP”) and amygdala (“Amy”). **D,E.** This pattern continues into P4 and adulthood (not shown). E is more caudal level than D. Expression in the hippocampus is strongest in CA3 and hilus, whereas multiple nuclei in amygdala show strong expression of *Cyp26b1* (E). Scale bars, A,B: 500μm, C-E:1mm.

**Fig.S2 Time-course of *Sst* expression in medial PFC of postnatal mice. A.** Left is to the midline. *Sst* mRNA was detected by *in situ* hybridization using a Tyramide Signal Amplification (TSA) system, followed by immunostaining with anti-SOX6 antibody. **B.**The ratio of *Sst*+; SOX6+ cells to SOX6+ cells reached a plateau value of ∼0.4 by P7. At P0, a much smaller portion of SOX6+ cells expressed *Sst* mRNA. Each dot indicates an average number of cells in medial PFC of at least 3 sections of a wildtype brain.

**Fig.S3 Recombination in *Synaptotagmin6-Cre* (*Syt6-Cre*) transgene mice**

Expression of ZSGreen in *Syt6-Cre/+*; *Ai6* (ZSGreen Cre reporter) mice at E12.5 (A), E14.5 (B) and P0 (C-E). All sections are coronal and the midline is to the left. At E12.5, expression of ZSGreen reporter is found in meninges (A, arrow) and preplate (A, arrowhead), but not in rest of the cortex or medial and lateral ganglionic eminences (MGE and LGE). At E14.5, a small number of cortical cells (B, double arrows) below the marginal zone (B, arrowhead) start to express ZSGreen. At P0, many layer 6 cells of frontal cortex express ZSGreen (C, arrow), but not in more caudal neocortex (D, “Ncx”), CA1, CA3 and hilus regions of the hippocampus (E, note that strong signal is found in the meninges of the hippocampus) or the amygdala (F, “Amy”). ic, internal capsule. Scale bar, 200μm

**Fig.S4 Generation of *Cyp26b1* CKO mice**

In situ hybridization of frontal sections of P8 (A,B,D-F, G,H, J-L) or P14 (C,I) *Cyp26b1* conditional knockout (G-L) and control littermate (A-F) brains. *Cyp26b1* was conditionally knocked out using the *Syt6-Cre* transgene. *Syt6* is expressed in layer 6 of both control (A) and *Cyp26b1* knockout (G) brain (arrow). Expression of *Cyp26b1* in layer 6 of frontal cortex (arrow) is detected in control, but not in *Cyp26b1* knockout brain at P8 (B,H) and P14 (C,I). Expression of *Cyp26b1* in agranular insula is unchanged in *Cyp26b1* knockouts (arrowhead in B,C,H,I). Expression of *Cyp26b1* in CA3 and hilus region of the hippocampus (D,J, arrowhead), globus pallidus (E,K, arrow) and amygdala (F,L, arrowhead) is unchanged in *Cyp26b1* knockouts. Scale bar, 1mm.

**Fig.S5 No significant changes in the number of *Sst-*, *Vip-* and *Lhx6-*expressing interneurons in medial PFC of *Cyp26b1* knockout mice**

**A-F.** In situ hybridization of frontal sections of P14 *Cyp26b1* conditional knockout mice (D-F) and littermate controls (A-C). Expression of *Somatostatin (Sst)* (A,D), *Vip* (B,E) and *Lhx6* (C,F) is shown. Binning and cell counts were done in the same was as shown in Fig.4 for *Pvalb*-expressing cells. Scale bar, 1mm. **G-I.** Result of statistical analysis. Each line connecting red and blue dots represents a pair of brains analyzed in the same experiment (n=5). P-values of paired ratio t-test for individual layer are shown. **J,K.** At P21, density of *Pvalb*-expressing cells (J), but not *Sst*-expressing cells, is increased in *Cyp26b1* conditional knockout mice. In repeated measures two-way ANOVA, the pvalues for layer (superficial vs deep), pair (between control and knockout), and interactions (between layer and pair) are 0.0047, 0.0287 and 0.0637 (*Pvalb*); 0.0065, 0.3621 and 0.7609 (*Sst*), **L-N.** No significant changes in the density of *Pvalb*-, *Sst-* and *Vip-*expressing interneurons in medial PFC of adult (P56-P67) *Cyp26b1* knockout mice. Each line connecting red and blue dots represents a pair of brains analyzed in the same experiment (n=4). P-values of paired t-test for individual layer are shown.

**Fig.S6: Thalamus-PFC disconnection in *Gbx2* mutant mice**

**A, B.** NetrinG1 immunostaining at E16.5. In control mice, NetrinG1-labeled thalamocortical axons are visible in coronal sections of frontal cortex. Arrowhead in A shows the medial PFC, where robust labeling is detected. In contrast, NetrinG1-labeling is barely detectable in the frontal cortex of *Gbx2* mutant mice, including the medial PFC (arrowhead in B). scale bar, 200μm.

**C-F.** DiI labeling at P14. DiI placement in medial PFC retrogradely labels medial thalamic nuclei (show more details) in the control brains (**C,D**). In *Gbx2* mutants, the label is severely reduced (**E,F**), indicating the deficiency of both thalamocortical and corticothalamic projections.

**Fig.S7 Expression of *RORβ* and *Lmo4* is qualitatively normal in the PFC of *Gbx2* mutant mice at postnatal day 8 (P8).**

**A,B.** Expression of *RORβ* in layer 4 is comparable between control (A) and *Gbx2* mutant (cko) mice (B)(arrows). **C,D.** Laminar expression patterns of *Lmo4* also appear unchanged in *Gbx2* mutants. scale bar=1mm.

